# A quick and reliable menthol-induced bleaching protocol for the Caribbean staghorn coral, *Acropora cervicornis*

**DOI:** 10.1101/2025.09.29.679371

**Authors:** J. Grace Klinges, Marina Villoch Diaz-Mauriño, Roger M. Wilder, Maia C. Erbes, Erinn M. Muller, Cory J. Krediet

**Author notes:** Corresponding Author: J. Grace Klinges.

## Abstract

Corals and dinoflagellate algae form a unique mutualistic symbiosis that provides the energetic and structural foundation for shallow coral reef ecosystems. Despite the long success of this partnership in oligotrophic seas, coral reefs are in decline due to increasing threats from rising seawater temperatures and disease, both of which can lead to bleaching and mortality. In order to better understand the mechanisms that underpin this mutualism, it may be necessary to dismantle the coral-algal symbiosis. Previous studies have experimentally bleached corals using thermal stress, photosynthetic inhibitors (DCMU), and menthol. We compared lab-induced bleaching of staghorn coral *Acropora cervicornis* by menthol treatment to traditional thermal stress. The larger aim was to adapt existing bleaching protocols to this important coral species used in restoration as a guide for future studies. Bleaching in corals treated with menthol or exposed to elevated temperature stress (31°C) was monitored by measuring photosynthetic activity determined by Fv/Fm using pulse-amplitude modulated (PAM) fluorescence. Corals were also monitored for symbiont density and overall health using the CoralWatch Coral Health Chart card throughout the experiment. We found that *A. cervicornis* bleached in response to both menthol treatment and thermal stress, but menthol treatment was more effective at reducing algal symbiont photosynthetic capacity (Fv/Fm) without negatively affecting the health of the coral. Our results indicate that menthol treatment at 0.38 mM rendered staghorn coral aposymbiotic within fourteen days without any visual or physiological damage to the coral. This study provides a simple and effective menthol-bleaching treatment protocol for future studies on staghorn coral.

## INTRODUCTION

The co-evolved mutualism between corals and their photosynthetic algal symbionts (family Symbiodiniaceae) provides the foundation for shallow coral reefs in oligotrophic waters. However, one of the greatest threats to coral-reef ecosystems in the present era of global climate change is the increasing frequency of coral-bleaching events (Brown, 1997; Douglas, 2003; Lesser, 2007; Hoegh-Guldberg et al., 2017). These bleaching events are often associated with high sea-surface temperatures (Douglas, 2003; Weis, 2008). Because the coral animals rely on photosynthetic products from their algal symbionts for much of their metabolic requirements, the breakdown of the symbiosis can result in coral death (Davy et al., 2012; Helgoe et al., 2024).

The ability of corals to switch algal symbiont communities after natural bleaching events has been well-documented (Cunning et al., 2015; Boulotte et al., 2016; Quigley et al., 2022). Recently, researchers have attempted to experimentally manipulate algal symbiont communities in *Acropora* by introducing or enriching thermotolerant strains of Symbiodiniaceae to enhance coral bleaching resistance (Matsuda et al., 2022; Buzzoni et al., 2023). These efforts include inoculation with stress-tolerant symbionts at various coral life stages, though the persistence of thermally-tolerant symbiont lineages outside of nursery contexts has been inconsistent (Abrego et al., 2009; Karp et al., 2025). In order to better understand the mechanisms of bleaching resistance in corals, it may useful to experimentally separate the members of the holobiont (the coral and all of its associated microorganisms, including endosymbiotic dinoflagellate of the family Symbiodiniaceae). Further, as cryopreservation is increasingly employed to preserve coral lineages, the most effective preservation methods require bleaching corals prior to vitrification due to the differing cryo-permeabilities of algal symbionts and coral tissue (Hagedorn et al., 2010, 2013; Lager et al., 2023). To functionally disassemble the coral-algal symbiosis, corals can be bleached using temperature stress (either cold shock or elevated temperature, Steen and Muscatine, 1987; Muscatine et al., 1991; Fitt et al., 2001) or photosynthetic inhibitors (e.g. DCMU, Jones, 2004; Wang et al., 2012; Matthews et al., 2016; Bauer et al., 2025). However, these traditional methods to bleach corals have also led to mortality in corals and sea anemones (Watanabe et al., 2006; Eakin et al., 2010; Muller et al., 2018).

Based on successful trials of menthol in inducing bleaching in other species of coral and anemones, we elected to compare the use of menthol with more traditional methods (i.e., thermal stress) of inducing bleaching in *Acropora cervicornis* in an effort to decrease mortality and coral stress in this important restoration species. Menthol, a compound found in mint-related plants, has been studied for its ability to trigger stress responses in coral similar to those caused by environmental stressors like temperature fluctuations or light stress (Wang et al., 2012; Matthews et al., 2016). Menthol has primarily been used in invertebrate systems for its anesthetic properties (Moore, 1989; Janes, 2008; Winlow et al., 2018; Yamazaki et al., 2024) but recent studies have demonstrated its ability to inhibit photosynthesis (Wang et al., 2017; Clowez et al., 2021) and cause bleaching in corals and anemones (Wang et al., 2012; Matthews et al., 2016; Puntin et al., 2024; Bauer et al., 2025). By exposing corals to menthol, we aimed to simulate bleaching events in a controlled environment and to develop an efficient bleaching protocol for *Acropora cervicornis*, allowing us to study the physiological and molecular responses of corals to bleaching without subjecting them to actual thermal stress. This method has shown promise in aiding our understanding of coral bleaching mechanisms and potential mitigation strategies.

## MATERIALS AND METHODS

All experiments were conducted at Mote Marine Laboratory’s International Center for Coral Reef Research in Summerland Key, Florida (24° 39′ 41.9′′ N, 81° 27′ 15.5′′ W). Replicate fragments of the staghorn coral *Acropora cervicornis* genotype ML-31 were collected from Mote Marine Laboratory’s *in situ* nursery in April 2023, and were fragmented into 5 cm-long ramets and mounted to ceramic plugs (Boston Aqua Farms, Windham, NH). Fragments were allowed to acclimate to *ex situ* conditions for one month prior to experimentation. Corals were housed in 5-gallon aquaria held in temperature-controlled, flow-through seawater raceways (∼170 gallons) with natural locally sourced sea water from the Atlantic side of the Keys. Sand- and particle-filtered water was fed from header tanks to aquaria at a flow rate of 7.2 L/hr. Aquaria were located outdoors under natural light regimes with the addition of 75% shade cloth from 11:30 am to 2:30 pm daily to maintain PAR at ∼300 μM/s during daylight hours. Five replicate corals were placed in each aquarium, and the egg crate racks holding corals were set to be 18 cm below the surface of the water. Aquaria were divided between two flow-through seawater raceways (20 aquaria per raceway), which allowed for temperature regulation of individual aquaria. Raceway water was prevented from entering aquaria by maintaining water levels below in and outflow holes using a standpipe. All corals were fed a standard diet of MicroVore, Zooplanktos-S, and Reef Snow (all Brightwell Aquatics, Fort Payne, AL) twice per week.

For comparison to menthol treatments, 15 coral fragments were bleached using elevated temperature (31.5°C) and an additional subset was maintained at ambient temperature (27.5°C) with no menthol treatment as a control. Elevated temperature treatments were performed in 5-gallon aquaria within outdoor raceways under ambient lighting. Temperature was controlled by a boiler and chiller using a dual heat exchanger system connected to header tanks and individual raceways. Header tank pH was stabilized at ∼8.0 by aeration and mixed via a venturi pump system. Elevated temperatures were achieved by incrementally increasing raceway temperature by 0.5 °C per day for eight days to reach 31.5 °C. Temperature, pH, dissolved oxygen, and salinity were measured in all raceways using a YSI (YSI Pro Plus, Xylem Inc, Washington, DC, USA) twice a day (8:00am and 12:00pm) three days a week (Monday, Wednesday, and Friday). Menthol incubation was performed on a total of 15 fragments for 6 hrs (∼11 am to 5 pm) in aerated aquaria filled with 17L of 0.2 μm filtered seawater. Aquaria were maintained indoors in low-light conditions with ambient light provided from a nearby window (∼100 PAR as measured by Licor Li-1500 and Li-192 underwater quantum sensor) (Lager et al., 2023, 2024). These conditions were selected to evaluate response to menthol alone, as light alone (350 PAR) has been shown to bleach corals (Lager et al., 2024). Temperatures were maintained at 27°C using 100W titanium aquarium heaters (Bulk Reef Supply) with Helio temperature controllers and tanks were aerated with an aquarium air pump. Additional water movement was provided using a 30 gph aquarium water pump. Three doses of menthol were tested, as in Wang et al. (2012): 0.19 mM, 0.38 mM, and 0.58 mM (99%, Sigma Aldrich, St. Louis, MO). Menthol was added to aquaria as 20% menthol in ethanol (w/v) (N = 5 fragments per dose treatment). Menthol in ethanol was added prior to introduction of corals to allow complete dissolution. Aquarium water and menthol were replaced every other day to maintain water clarity as symbionts were ejected. After incubation, coral fragments were returned to the holding raceway and held at 27°C and ambient outdoor light for 17-18 h (overnight) under conditions identical to the control treatment. The fragments were cycled daily between the menthol bath and raceway conditions for two weeks until they were fully bleached (reached the lowest color ranking on a CoralWatch Coral Health Chart, (Siebeck et al., 2008). After two weeks, corals were allowed to recover in raceways for one month to observe continuing impacts of treatment on coral health.

To monitor bleaching progression throughout the menthol or thermal stress treatments, we measured photosynthetic activity as Fv/Fm through pulse-amplitude modulated (PAM) fluorometry using a Junior-PAM (Heinz Walz, Germany). The Junior-PAM employs a blue LED (pulsed through a fiber optic cable) for pulse modulated excitation light, actinic illumination, and saturation pulse analysis of photosystem II. The fluorometer has a far-red LED for selective excitation of photosystem II, which is needed for determination of F0’ (baseline) fluorescence. Photosynthetic activity was measured at day 0, 7, and 14 of the menthol treatments and at day 0, 7, 14, and 28 of the elevated temperature treatment. Prior to measurement, corals were dark-adapted for 30 minutes to capture maximum photochemical efficiency of photosystem II (as Fv/Fm). Measuring light intensity was set to 6, SAT-Pulse Intensity was set to 12, SAT-Pulse width was set to 0.6, and Gain was set to 3. Measurements were taken as three saturation pulses back-to-back in triplicate per coral, selecting different areas of the coral fragment to acquire each measurement. F_v_/F_m_ was calculated from measured minimum and maximum fluorescence (F_o_ and F_m_) as (F_m_ - F_o_)/F_m_. Data were averaged within each replicate fragment (3 measurements per fragment). Differences in Fv/Fm across timepoints were assessed using a Kruskal-Wallis rank sum test, followed by pairwise Wilcoxon rank sum tests with false discovery rate (FDR) correction for multiple comparisons. Rough assessments of algal symbiont concentrations and coral health were made weekly using a CoralWatch Coral Health Chart (Siebeck et al., 2008). The CoralWatch Coral Health Chart provides a six-point scale with which changes in coral color can be measured as an indicator of symbiont density. Photographs of each coral individual were taken contemporaneously with coral health card measurements.

## RESULTS AND DISCUSSION

Throughout the experiment, corals exhibited no necrosis across any dose of menthol-treated corals, with corals continuing to feed and exhibiting good polyp extension for the first week of menthol exposure (Figure 1). Limited to no polyp extension was observed in corals bleached with elevated temperature (31°C), and tissue necrosis was observed on these corals (Supp. Figure 1). We observed visible differences in algal symbiont density between menthol-treated corals and untreated controls as early as 48 hours into treatment (Figure 1, Figure 2, Supp. Figure 5). On the third day of treatment, corals dosed with 0.58 mM menthol exhibited retracted tissue and significant mesenterial filament extension (Supp. Figure 2). Corals dosed with 0.38 mM had moderate mesenterial filament extension, while corals dosed with 0.19 mM menthol displayed no mesenterial filaments and polyps were extended (Figure 1).

**Figure 1.**
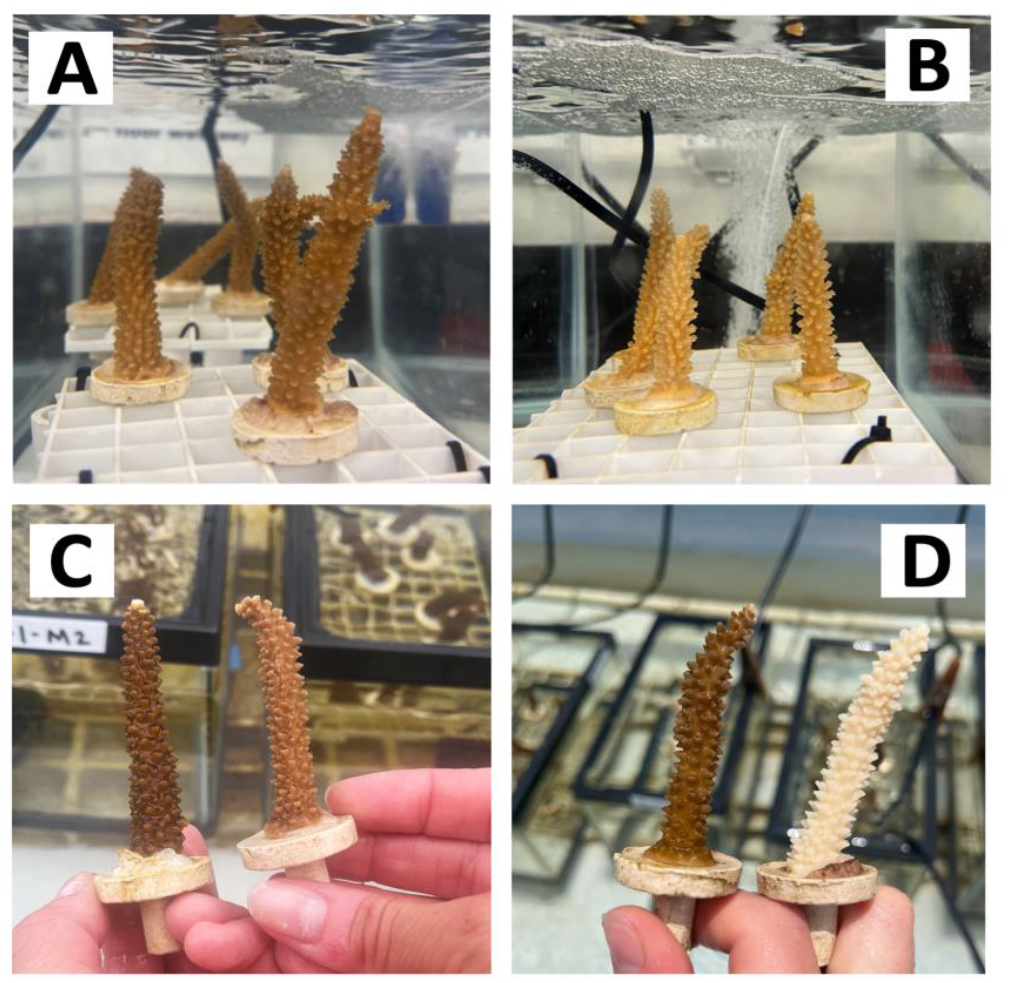
Corals exhibited polyp extension, indicative of good health during menthol bleaching. A) Corals on the first day of menthol treatment (0.38 mM dose). B) Corals on the third day of menthol treatment (0.38 mM dose). C) Untreated coral (left) compared to menthol-treated coral (0.58 mM dose) at 48 hours. D) Untreated coral (left) compared to menthol-treated coral (0.58 mM dose) at 2 weeks.

**Figure 2.**
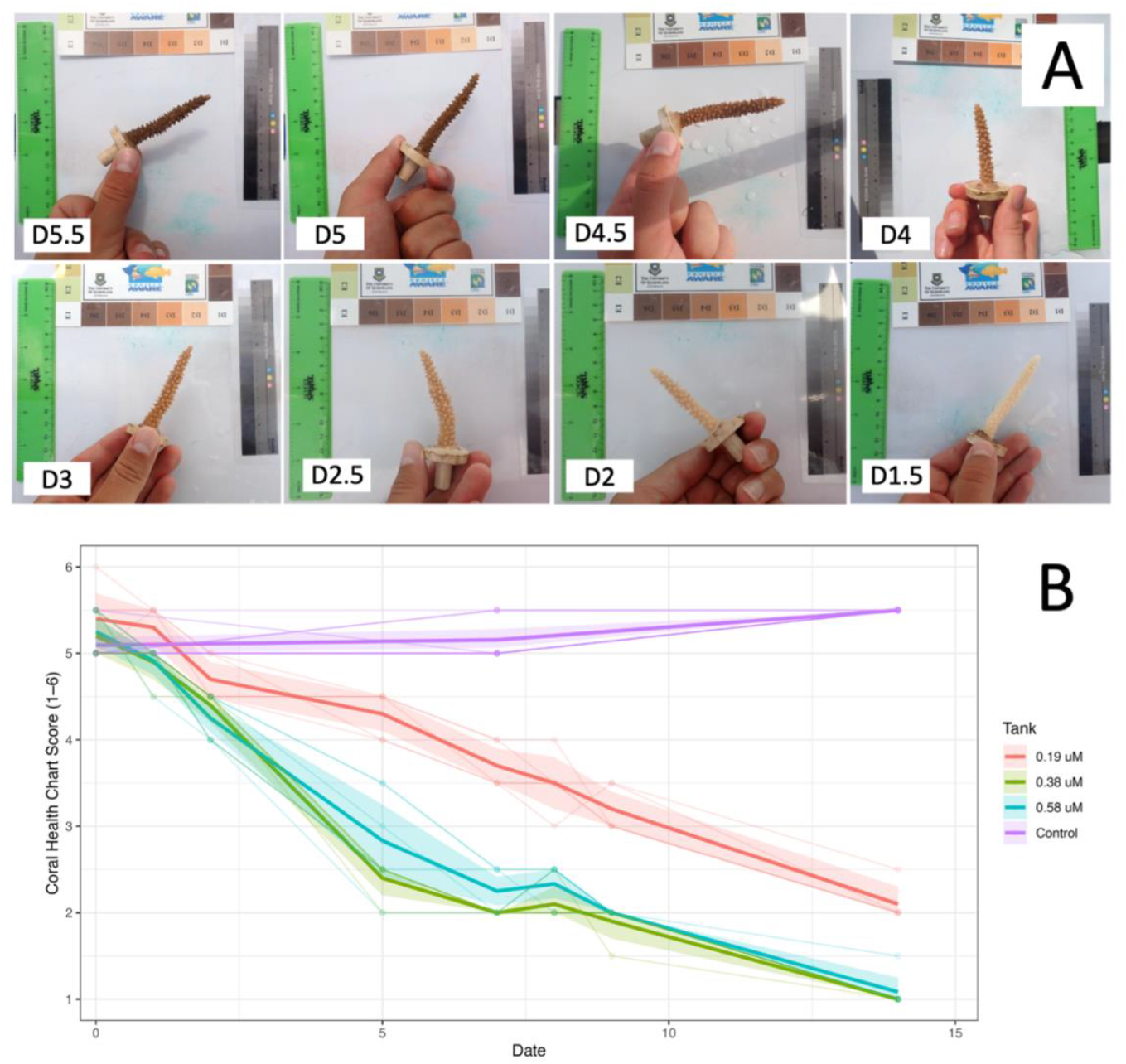
**A**. The CoralWatch Coral Health Chart was used to estimate coral health scores using a scale from D1-D6. Health scores (1–6) were assigned by visual comparison to a standardized color reference card to ensure consistent scoring across observers and timepoints. Representative images for various scores are shown. B. Mean health scores (±95% confidence intervals) of genotypes across treatments over time. Points represent observed mean 184 health scores (as determined from CoralWatch Coral Health Chart) for each tank treatment at each sampling date, calculated across all genotypes within a treatment. Error ribbons denote 95% confidence intervals around the treatment means. Individual genotype-level observations (faint lines and points) are shown to illustrate within treatment variability. See Supplemental Figure 5 for smoothed linear regression.

Measurements of photosynthetic efficiency as Fv/Fm were taken for all corals through pulse-amplitude modulated (PAM) fluorometry using a Junior-PAM prior to treatment and at day 7 and day 14 of menthol treatment (Figure 3) and at day 0, day 7, day 21, and day 28 for corals bleached through elevated temperature (Figure 4, Supp. Figure 3). High temperature (31°C) was effective at bleaching corals by three weeks of exposure (Figure 4, Supp. Table 1). The interaction of treatment and timepoint had a significant impact on photosynthetic efficiency of menthol-treated corals as measured by PAM (chi-squared = 42.271, df = 8, p-value = 1.205e-06). All three doses of menthol, as well as high temperature, were effective in reducing photosynthetic activity from initial/control values of around 0.65 (Supp. Table 2), however, bleaching was much more effective at higher doses (0.38 and 0.58 mM) of menthol after only seven days of treatment (Figure 3, Supp. Figure 4, Supp. Table 2). Although Fv/Fm values of corals treated with 0.19 mM menthol were significantly lower than initial values by as early as one week (p < 0.5, Supp. Table 2), corals treated with either 0.38 or 0.58 mM menthol had significantly lower Fv/Fm values than both initial values and the 0.19 mM dose (p < 0.5, Supp. Table 2, Figure 3). Indeed, average Fv/Fm values were significantly lower in menthol-treated corals after only seven days compared to high temperature-treated corals after 28 days (Figure 4, Supp. Table 3). As observed in other studies (Wang et al., 2012, Bauer et al., 2025), we found that the highest dose test (0.58 mM menthol) induced partial fragment mortality that was observed after the two-week treatment period, as corals recovered from treatment. The lowest dose test (0.19 mM menthol) significantly reduced Fv/Fm from baseline measurements and compared to untreated controls (p < 0.05, Supp. Table 2), but Fv/Fm at 2 weeks for 0.19 mM menthol was significantly higher than Fv/Fm from both higher menthol doses and from high temperature-bleached corals (p < 0.05, Supp. Figure 4, Supp Table 3).

**Figure 3.**
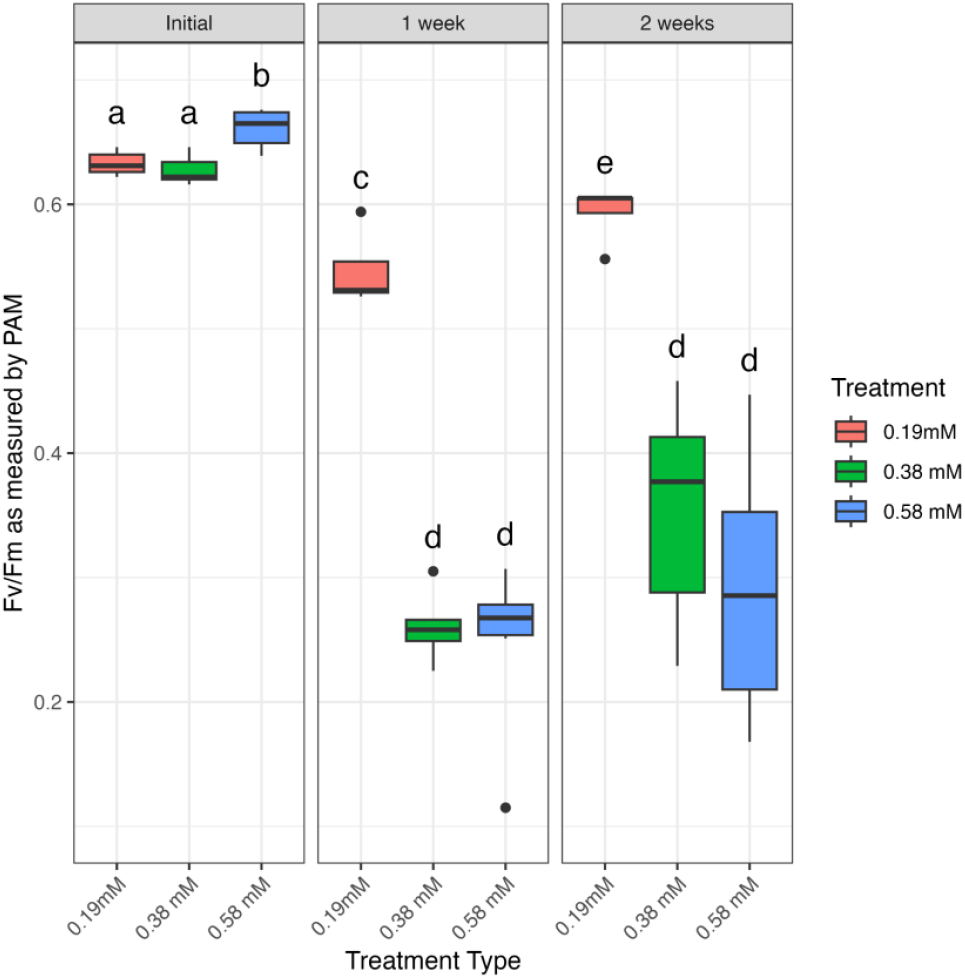
Fv/Fm as measured by Junior-PAM, compared between three doses of menthol (0.19 mM, 0.38 mM, and 213 0.58 mM) prior to treatment and after one and two weeks of menthol treatment. Boxes that share a letter are not significantly different from each other at α = 0.05 (Kruskal-Wallis with Pairwise Wilcoxon).

We concluded that menthol treatment is highly effective at bleaching *A. cervicornis* without significant impacts on coral health and selected 0.38 mM as the optimal dose for this species. Corals treated with 0.38 mM and 0.58 mM menthol showed similar reduction in both Fv/Fm and visible symbiont density (Figure 2, Figure 3), and resulted in a lower Fv/Fm than thermally-bleached corals after only one week (Figure 4). As treatment with 0.38 mM menthol was sufficient to achieve bleaching, and as this treatment with did not cause mortality as has been reported previously (Wang et al. 2012), we suggest this dose be used for bleaching and cryopreservation studies with this species moving forward.

**Figure 4.**
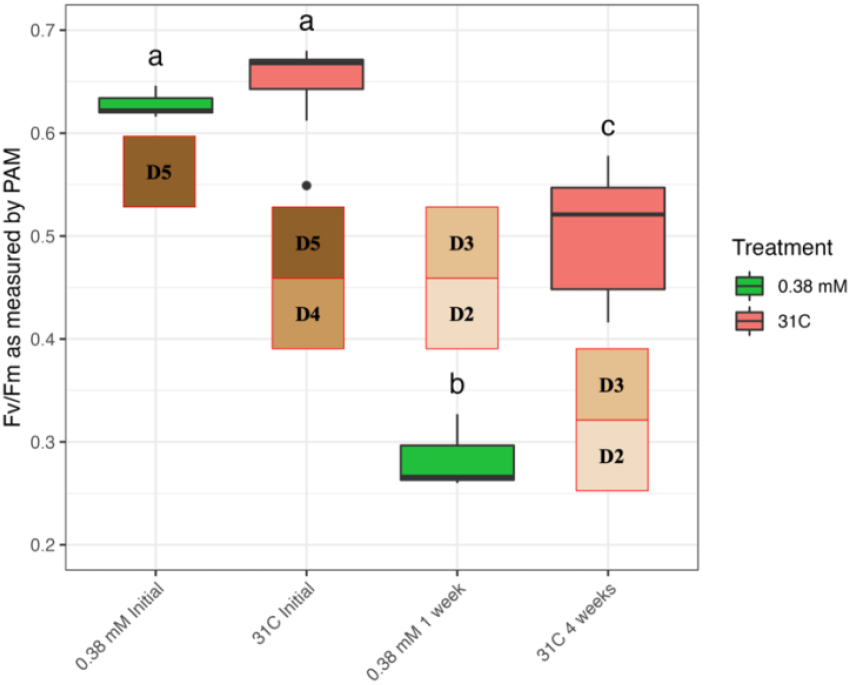
Fv/Fm as measured by Junior-PAM, comparing corals bleached using 0.38 mM menthol to corals bleached using the more traditional method of thermal stress at 31°C at day 0 and at the end of treatment (7 days for menthol and 4 weeks for thermal bleaching). Boxes that share a letter are not significantly different from each other at α = 0.05 (Kruskal-Wallis with Pairwise Wilcoxon). Also shown for each timepoint is the corresponding average coral health score, acquired using CoralWatch Coral Health Chart.

Our results demonstrate that menthol treatment is an effective and reliable method to induce bleaching in staghorn coral, *Acropora cervicornis*, compared to traditional methods using elevated temperature. Menthol treatment yielded significant reduction in algal symbiont densities and significantly reduced Fv/Fm values without compromising the integrity of the coral host. Interestingly, 1 week of menthol bleaching reduced Fv/Fm values further than 4 weeks of temperature bleaching, though both treatments had similar algal symbiont density as visually assessed by CoralWatch card. Temperature bleaching induced total or partial mortality in 93.7% of frags that continued to develop after temperature stress alleviated. Although no mortality was seen during menthol treatment, partial mortality occurred in subsequent weeks after the experiment, impacting 56.3% of frags and which was seemingly independent of treatment. As corals were not re-inoculated with algal symbionts after treatment, this mortality may be related to reduced nutrition from loss of symbionts rather than directly related to treatment. As one week of menthol treatment was sufficient to greatly reduce photosynthetic efficiency, though not to fully bleach corals, a treatment duration between one and two weeks could be sufficient for applications such as cryopreservation that require high survival rates after bleaching. Enhanced methods for recovery of algal symbionts after bleaching have been developed that could further reduce mortality rates after treatment (Morgans et al., 2020; Scharfenstein et al., 2022).

As the field of coral restoration continues to build and provide insight into the mechanisms of thermal tolerance and disease resilience, there may be a need to experimentally bleach corals to form aposymbiotic individuals. This has already proven valuable in the coral model system, *Aiptasia* (Goulet et al., 2005; Lehnert et al., 2012; Dungan et al., 2020; Roberty et al., 2024) as well as for the cryopreservation of coral microfragments (Lager et al., 2023, 2024). The ability to render and keep corals in the aposymbiotic state may allow for investigation of symbiont establishment and maintenance, dynamics of host-symbiont recognition, symbiosis function in response to climate change, manipulation of algal symbionts and host microbiome, application of beneficial microorganisms of coral (BMCs), cryopreservation of full coral fragments, and mechanisms of coral bleaching.

## AUTHOR CONTRIBUTIONS

J.G.K., C.J.K., AND E.M.M. conceptualized the research. J.G.K. and C.J.K. designed the experiment. J.G.K., M.V.D.M., R.M.M, and M.C.E. carried out the experiment. J.G.K. analyzed data. J.G.K. and C.J.K. interpreted data. J.G.K. and C.J.K. wrote the manuscript and all authors contributed edits to the manuscript.

## ACKNOWLEDGMENTS

This work was funded by the Eppley Foundation for Research and the Mote Marine Laboratory Protect Our Reefs Grant with award number POR 2021-9. We thank staff at Mote Marine Laboratory’s Elizabeth Moore International Center for Coral Reef Research and Restoration for expertise and logistical help including Eleftherios Karabelas, Kyle Knoblock, and Zachary Craig. The use of nursery-grown corals was authorized by the Florida Keys National Marine Sanctuary under Special Activity License SAL-21-2048A and SAL-222406A-SCRP.

## Supplementary Materials

### Supplemental Tables

**Supplemental Table 1.** Adjusted p-values from pairwise Wilcoxon rank sum tests comparing Fv/Fm values across timepoints for temperature-bleached corals (at 31°C) and untreated conspecifics across 4 weeks of treatment and one week of recovery. P-values were corrected for multiple comparisons using the false discovery rate (FDR) method. Significant p-values are bolded and in red.

|  | Control Initial | 31C Initial | 31C 1 week | Control 2 weeks | 31C 3 weeks | 31C 4 weeks | Control 4 weeks |
| --- | --- | --- | --- | --- | --- | --- | --- |
| 31C Initial | **0.0042** | NA | NA | NA | NA | NA | NA |
| 31C 1 week | **0.0144** | 0.4148 | NA | NA | NA | NA | NA |
| Control 2 weeks | 0.6061 | **0.0144** | 0.0584 | NA | NA | NA | NA |
| 31C 3 weeks | **2.09E-05** | **9.99E-06** | **9.99E-06** | **1.89E-05** | NA | 0.2826 | NA |
| 31C 4 weeks | **9.99E-06** | **9.99E-06** | **9.99E-06** | **9.99E-06** | 0.2826 | NA | NA |
| Control 4 weeks | 0.0759 | 0.0584 | 0.1864 | 0.2517 | **9.99E-06** | **9.99E-06** | NA |
| 31C 1 week recovery | **9.99E-06** | **9.99E-06** | **9.99E-06** | **9.99E-06** | 0.8209 | 0.3661 | **9.99E-06** |

**Supplemental Table 2.**
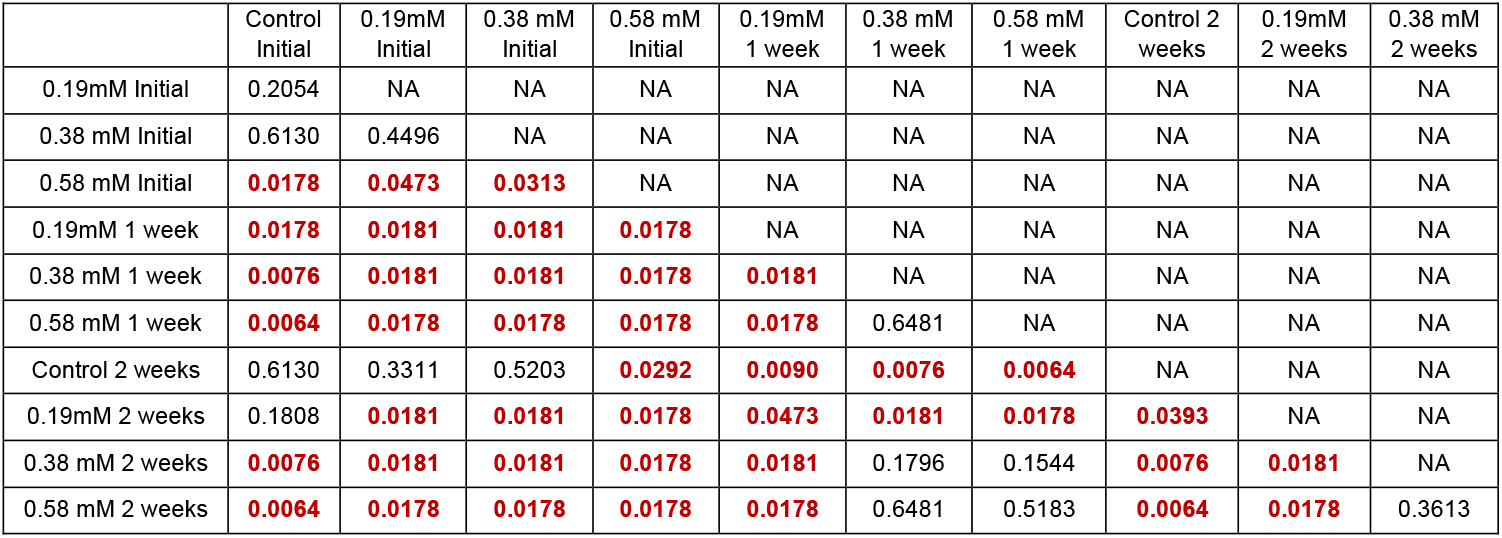
Adjusted p-values from pairwise Wilcoxon rank sum tests comparing Fv/Fm values across timepoints for menthol-bleached corals and untreated conspecifics at three doses across 2 weeks of treatment. P-values were corrected for multiple comparisons using the false discovery rate (FDR) method. Significant p-values are bolded and in red.

**Supplemental Table 3.** Adjusted p-values from pairwise Wilcoxon rank sum tests comparing Fv/Fm values across timepoints for menthol-bleached corals at three doses across 2 weeks of treatment and temperature-bleached corals (at 31°C) across 4 weeks of treatment. P-values were corrected for multiple comparisons using the false discovery rate (FDR) method. Significant p-values are bolded and in red.

|  | 0.19mM<br>Initial | 0.38<br>mM<br>Initial | 0.58<br>mM<br>Initial | 31C<br>Initial | 0.19mM<br>1 week | 0.38<br>mM 1<br>week | 0.58<br>mM 1<br>week | 31C 1<br>week | 0.19mM<br>2 weeks | 0.38<br>mM 2<br>weeks | 0.58<br>mM 2<br>weeks |
| --- | --- | --- | --- | --- | --- | --- | --- | --- | --- | --- | --- |
| 0.19mM<br>Initial | NA | NA | NA | NA | NA | NA | NA | NA | NA | NA | NA |
| 0.38 mM<br>Initial | 0.43336 | NA | NA | NA | NA | NA | NA | NA | NA | NA | NA |
| 0.58 mM<br>Initial | **0.04665** | **0.02987** | NA | NA | NA | NA | NA | NA | NA | NA | NA |
| 31C Initial | 0.05853 | 0.04892 | 1.00000 | NA | NA | NA | NA | NA | NA | NA | NA |
| 0.19mM 1<br>week | **0.01676** | **0.01676** | **0.01530** | **0.00713** | NA | NA | NA | NA | NA | NA | NA |
| 0.38 mM 1<br>week | **0.01676** | **0.01676** | **0.01530** | **0.00486** | **0.01676** | NA | NA | NA | NA | NA | NA |
| 0.58 mM 1<br>week | **0.01530** | **0.01530** | **0.01434** | **0.00380** | **0.01530** | 0.65805 | NA | NA | NA | NA | NA |
| 31C 1<br>week | 0.10836 | 0.06377 | 0.49020 | 0.42367 | **0.00890** | **0.00486** | **0.00380** | NA | NA | NA | NA |
| 0.19mM 2<br>weeks | **0.01676** | **0.01676** | **0.01530** | **0.01375** | **0.04678** | **0.01676** | **0.01530** | **0.01676** | NA | NA | NA |
| 0.38 mM 2<br>weeks | **0.01676** | **0.01676** | **0.01530** | **0.00486** | **0.01676** | 0.16636 | 0.14224 | **0.00486** | **0.01676** | NA | NA |
| 0.58 mM 2<br>weeks | **0.01530** | **0.01530** | **0.01434** | **0.00380** | **0.01530** | 0.65805 | 0.49361 | **0.00380** | **0.01530** | 0.35271 | NA |
| 31C 4<br>weeks | **0.00486** | **0.00486** | **0.00380** | **0.00011** | 0.19645 | **0.00486** | **0.00380** | **0.00011** | **0.00713** | **0.01375** | **0.00552** |

### Supplemental Figures

**Supplemental Figure 1.**
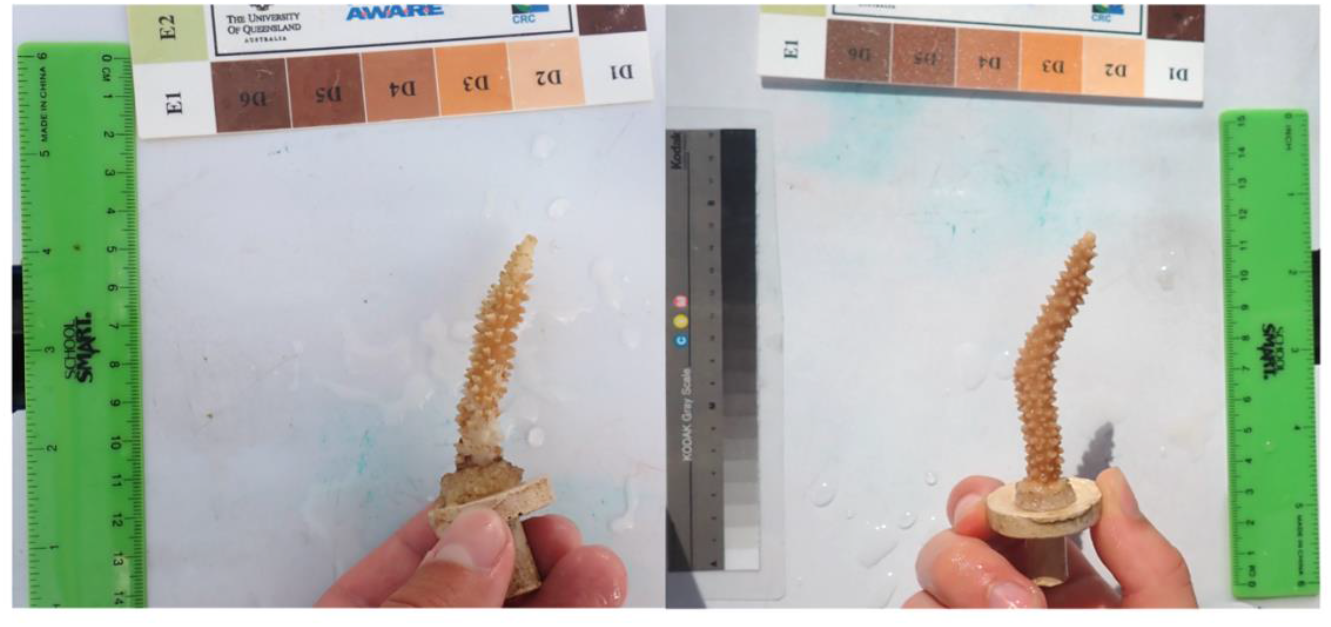
Coral bleached by high temperature indicating signs of partial mortality (left) compared to coral bleached by menthol with no signs of partial mortality. Both corals are bleached to a similar degree as measured by CoralWatch coral health chart.

**Supplemental Figure 2.**
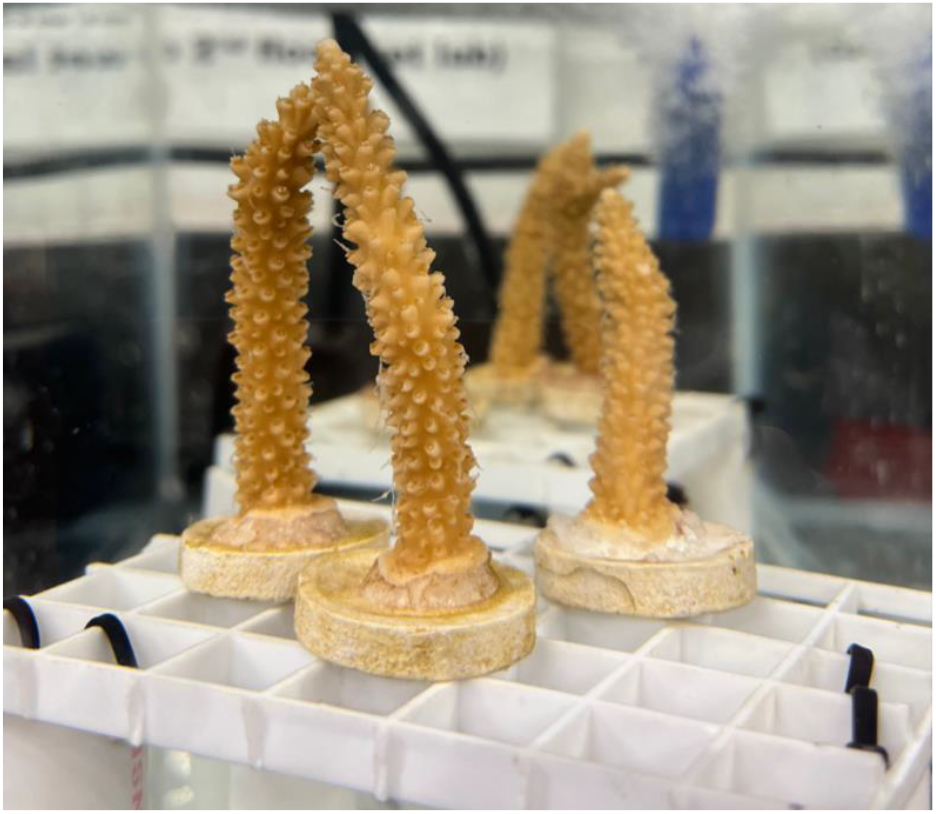
Mesenterial filament extension observed on menthol-treated corals.

**Supplemental Figure 3.**
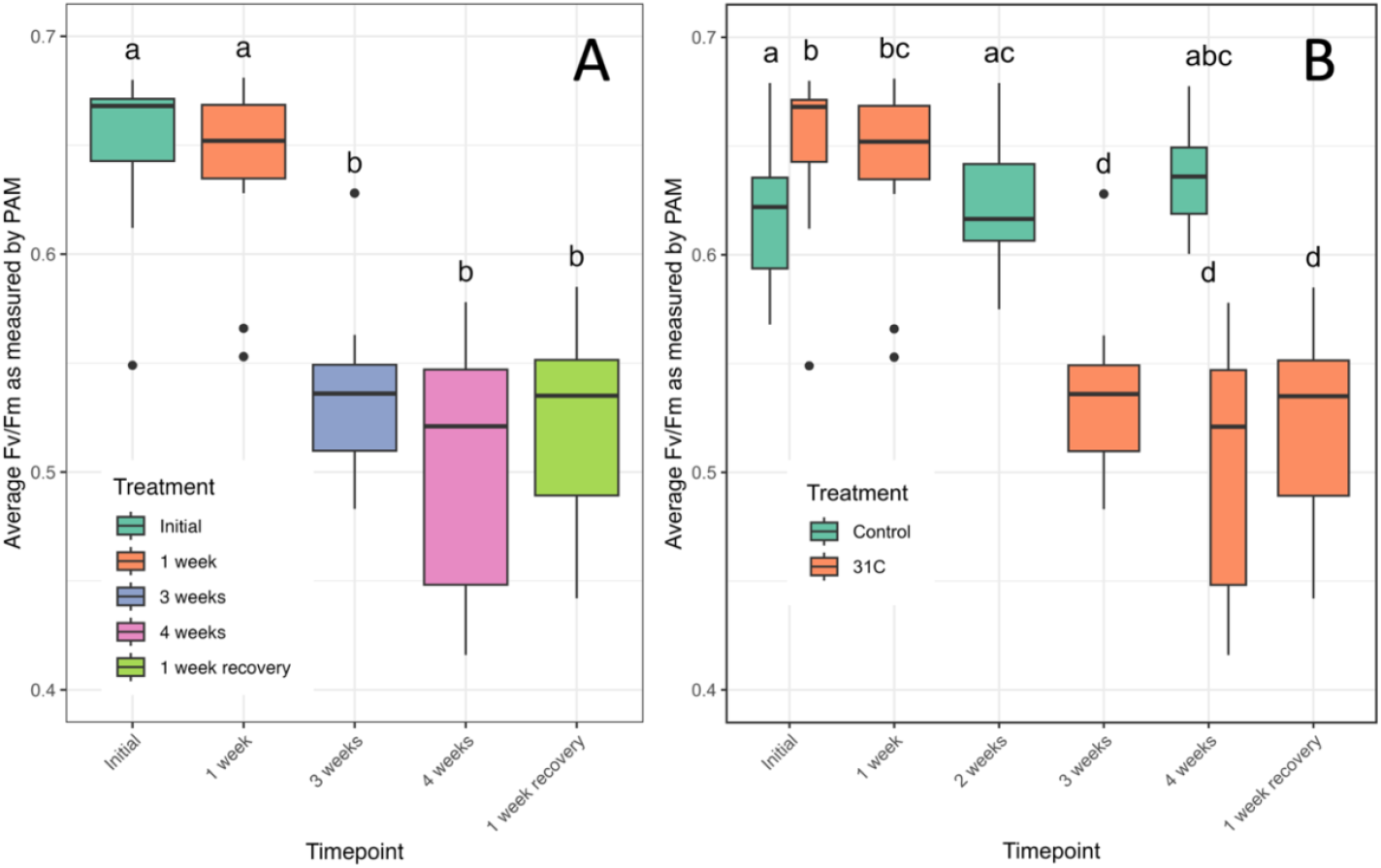
Fv/Fm as measured by Junior-PAM, shown for temperature-bleached corals (at 31°C) across 4 weeks of treatment and one week of recovery without controls (A) and with controls (B). Boxes that share a letter are not significantly different from each other at α= 0.05 (Kruskal-Wallis with Pairwise Wilcoxon).

**Supplemental Figure 4.**
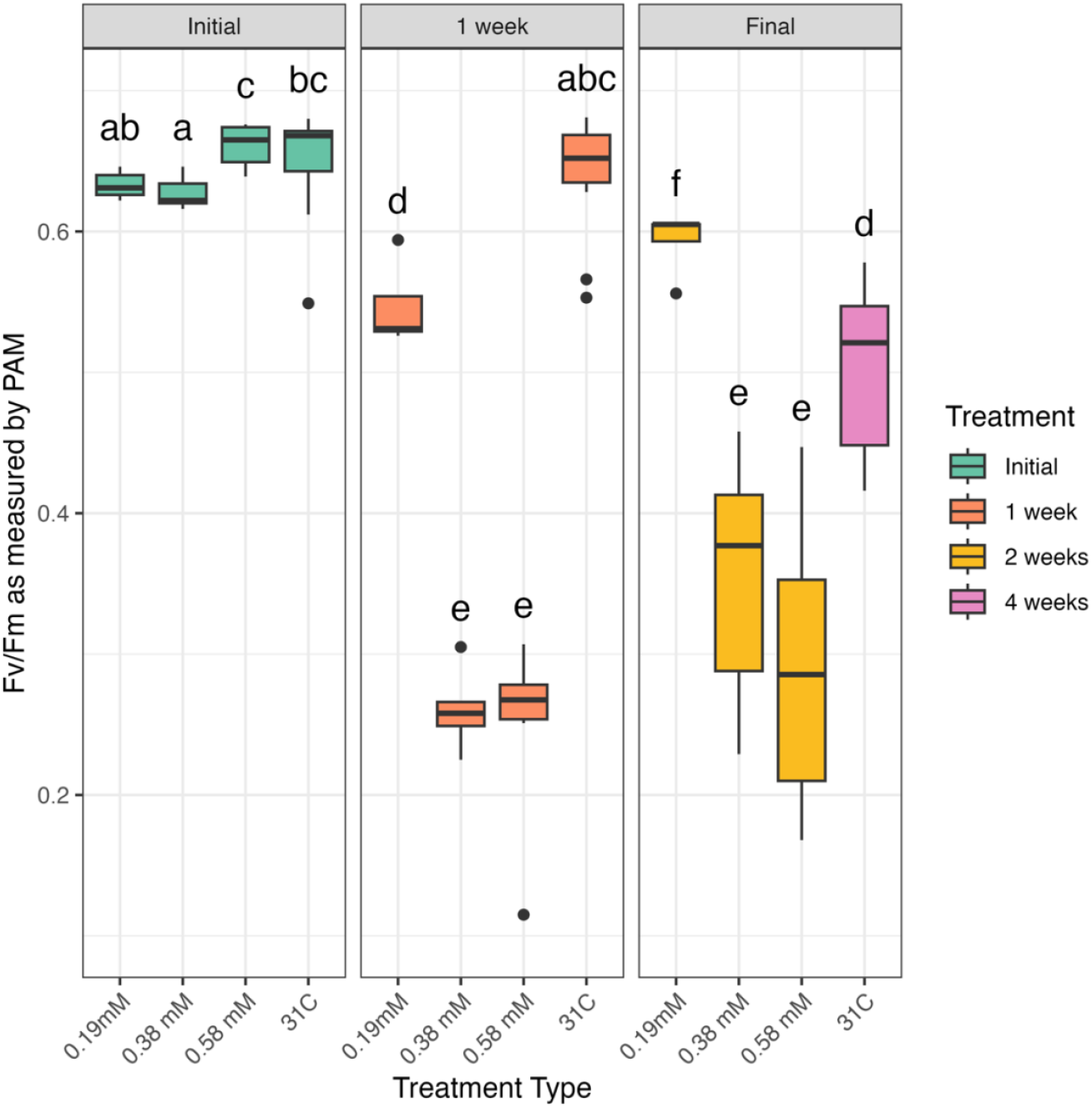
Fv/Fm as measured by Junior-PAM at initial, 1 week, and final timepoints for menthol- and temperature-bleached corals. The final timepoint for menthol-bleached corals was at 2 weeks and the final timepoint for temperature-bleached corals was at 4 weeks. Boxes that share a letter are not significantly different from each other at α= 0.05 (Kruskal-Wallis with Pairwise Wilcoxon).

**Supplemental Figure 5.**
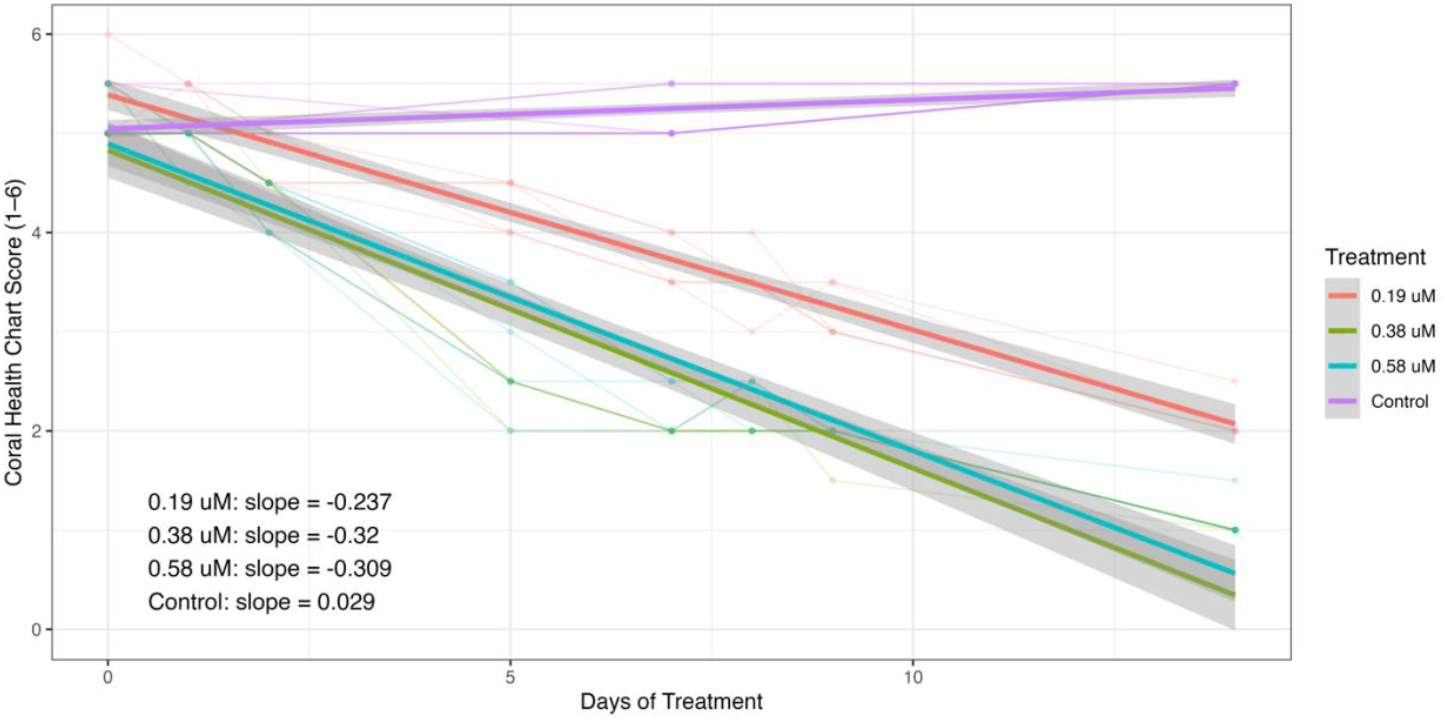
Health score trajectories of genotypes under different tank treatments over time, with slopes estimated from generalized linear models. Individual genotypes are shown as faint lines and points to illustrate within-treatment variability. Bold regression lines represent fitted slopes from a generalized linear model of health score as a function of time (days of treatment) and treatment. Reported slope values (lower left inset) indicate the rate of change in health score per day for each treatment. Shaded ribbons represent 95% confidence intervals around model predictions.

